# Grapevine phyllosphere pan-metagenomics reveals pan-microbiome structure, diversity, and functional roles in downy mildew resistance

**DOI:** 10.1101/2025.05.04.652149

**Authors:** Jingyun Jin, Xiangfeng Wang, Xuenan Zhang, Junjie Mei, Wei Zheng, Linling Guo, Haisheng Sun, Lili Zhang, Chonghuai Liu, Wenxiu Ye, Li Guo

## Abstract

Grapevines are among the most economically important fruit crops, and microbiome profoundly influence their health, yield, and quality. However, mechanistic insights into microbiome-orchestrated grapevine biology remain limited. Here, we conduct large-scale pan-metagenomic and pan-metatranscriptomic analyses of the phyllosphere microbiome from 107 grapevine accessions spanning 34 *Vitis* species. We show that the grapevine core microbiome is dominated by phylum Bacillota and Pseudomonadota. Leveraging PacBio sequencing, we assembled 19 high-quality metagenome-assembled genomes (MAGs) from the grapevine pan-microbiome, representing the first MAG reconstruction in plant-associated microbial communities using PacBio reads. These MAGs encode genes associated with antibiotic resistance, secondary metabolism, and carbohydrate-active enzymes (CAZymes), which could potentially influence grapevine biology. During downy mildew (DM) infection, DM-resistant grapevines exhibit significantly higher microbial network complexity than susceptible counterparts. Among the key taxa contributing to this complexity, Bacillota emerged as the dominant phylum with displaying strong abundance correlations with phylum Euglenozoa and Cyanobacteriota, and an isolated Bacillota species from the grapevine leaves, *Bacillus cereus*, demonstrated potent biocontrol activity against DM infection. Pan-metatranscriptomic analysis further revealed significant upregulation of eukaryotic microbial genes involved in primary and secondary metabolism. This pan-metagenomic study offers unprecedented insights into the complex structure, diversity and functional roles of grapevine phyllosphere microbiome, and presents valuable genomic and microbial resources for microbiome research and engineering to enhance viticulture productivity and quality.

## Introduction

Grapevines are perennial plants belonging to Vitaceae family renowned for their exceptional flavor as well as their significant economic and nutritional value. In 2023, global grape production reached 72 million tons, with China, Italy, France leading as the top producers, and the global production value of grapes was estimated at 92 billion USD [1]. In addition, grapes are rich in bioactive compounds, including polyphenols, phenolic acids, vitamins, and dietary fiber, which provide a variety of health benefits, such as anticancer, antidiabetic, cardioprotective, and gut-microbiome-regulating properties [2]. Despite remarkable advancements in grape breeding and cultivation in recent decades, grape production worldwide remains significantly threatened by diseases such as downy mildew (DM), powdery mildew, and gray mold, causing substantial economic losses across grape productions and related industries each year [3–7]. Especially, DM is the most destructive among the diseases that caused by the oomycete *Plasmopara viticola*, severely affecting grape production and quality by causing damages on leaves and fruit clusters [8]. With increasing evidence highlight the involvement of grapevine microbiome in DM infection [9,10], it is crucial to investigate grapevine-associated microbiome, which could serve as a foundation for developing environmentally friendly and sustainable microbiome-based biocontrol methods in viticulture.

Microbiome, often regarded as the second genome of plants, plays crucial roles in nutrient absorption, disease resistance, and overall plant health [4]. Microbiome profoundly affects grape productivity and the quality of grape-derived products such as wine and raisins [11–13]. The grapevine microbiomes primarily consist of bacteria and fungi, with Acidobacteria, Actinomycetota, Bacillota, Bacteroidota, Chloroflexi, and Pseudomonadota being the most prevalent phyla, based on previous 16s rRNA amplicon sequencing [4,9,14–16]. The microbiome composition varies across grapevine tissues with the highest microbial diversity observed in roots and gradually decreasing for trunks and leaves [4]. Studies show that grapevine genetic backgrounds and geographic origins are primary factors influencing their microbial compositions [11,14,17–24]. Despite the advancement, current plant microbiome studies primarily rely on 16S rRNA sequencing which yields limited insights into mechanistic understanding of microbiome functions compared to Illumina sequencing. However, Illumina sequencing falls short in assembling the complete microbial genomes given the limitation of short reads, impeding accurate annotations and functional analysis of metagenome. So far, long-read technologies mainly have been used in microbiome studies for human related samples [25,26], but remain underutilized in plant and agricultural metagenomics. In addition, metatranscriptomics, which crucial to understanding functional implications of microbiome by revealing microbiome gene expression changes, has not been widely adopted in plant microbiome research. Most importantly, integrating pan-metagenomics and pan-metatranscriptomics of plants to study the diversity of microbial profiles across plant species and cultivars have not been reported. These limitations hinder the exploration of grapevine microbiome and the application of microbial biocontrol in viticulture.

Here, we conducted a comprehensive pan-metagenomic analysis of grapevine leaves from 107 *Vitis* accessions (each accession corresponds to a unique *Vitis* cultivar, except for a few samples) spanning 34 species, utilizing Illumina, PacBio, RNA-seq and DM resistance phenome data. Through systematic profiling of the grapevine phyllosphere microbiome, we identified dominant microbial communities under pan-metagenomic framework. Leveraging PacBio metagenomic reads from the grapevine phyllosphere, we successfully generated 19 high-quality grapevine phyllospheric metagenome-assembled genomes (GPMAGs), and comprehensively annotated genes associated with antibiotic resistance, secondary metabolism, and CAZymes. Based on the phenome data, we revealed that the microbiome from DM-resistant grapevines exhibit significantly more complex microbial networks than susceptible ones. Additionally, we identified key genera potentially conferring resistance to DM infection, with Bacillota emerging as the dominant phylum exhibiting high abundance correlation with Euglenozoa and Cyanobacteriota, and experimentally validated the potent biocontrol activity of two *Bacillus cereus* strains against DM. Pan-metatranscriptomic analysis further highlighted significant changes in the expression of eukaryotic primary and secondary metabolism genes in response to DM infection. These findings provide novel insights into grapevine microbiome and valuable genomic and microbial resources for viticulture, laying a solid foundation for developing microbiome-based strategies to enhance grape health and promote sustainable production.

## Results

### Grapevine core microbiome is dominated by Bacillota and Pseudomonadota

To investigate the microbiome profile of the grapevine phyllosphere across grapevines, we collected a total of 107 leaf samples from 34 *Vitis* species covering 100 grapevine cultivars originated from various geographical locations with distinct genotypes and DM resistances [27] (**Fig. 1a, b and Table S1**), and conducted pan-metagenomic analysis as outlined in the workflow (**Fig. 1c**). The metagenomic DNA from the samples was sequenced using Illumina paired-end sequencing [27], yielding a total of 107 Illumina shotgun metagenomics datasets (**Fig. 1d**). The reads belonging to the grapevines were completely filtered out by mapping against a complete grapevine genome [27] (VHP-T2T.hap1, NCBI accession: GCA_046255155.1) (**Table S2**), with on average 4.4 million microbial reads remaining per sample. Taxonomic classification of these microbial reads using Kraken2 [28] identified a total of 1,326 genera across the grapevine phyllosphere (**Fig. 2a and Table S3**). Among these, Bacillota (43.6%) and Pseudomonadota (37.9%) dominated the microbiome profile at the phylum level, while *Lactiplantibacillus* (42.1%) and *Xanthomonas* (22.2%) were the two most abundant genera (**Fig. 2b**). Furthermore, kingdom-level profiling demonstrated that bacteria dominated the grapevine phyllosphere microbiome (91.3%), and other kingdoms accounted for less than 10% (**Fig. 2c**). Notably, the non-bacterial kingdoms exhibited dominating phyla, such as Euryarchaeota dominating Archaea, Ascomycota and Euglenozoa dominating Eukaryota, and Cressdnaviricota dominating viruses, reflecting specific phylum-level dominance for the grapevine microbiome. To examine worldwide representativeness of our microbial composition, we analyzed worldwide representative accessions from published Illumina data containing a global grapevine phyllosphere collection [29] using the same bioinformatic pipeline. The phylum-level microbial composition of the global microbiome generally consistent with our Illumina-based composition (**Fig. S1**), demonstrating the worldwide representativeness of our microbial composition. These results highlight the prevalence of Bacillota and Pseudomonadota and their potential roles in the grapevine microbiome.

**Fig. 1.**
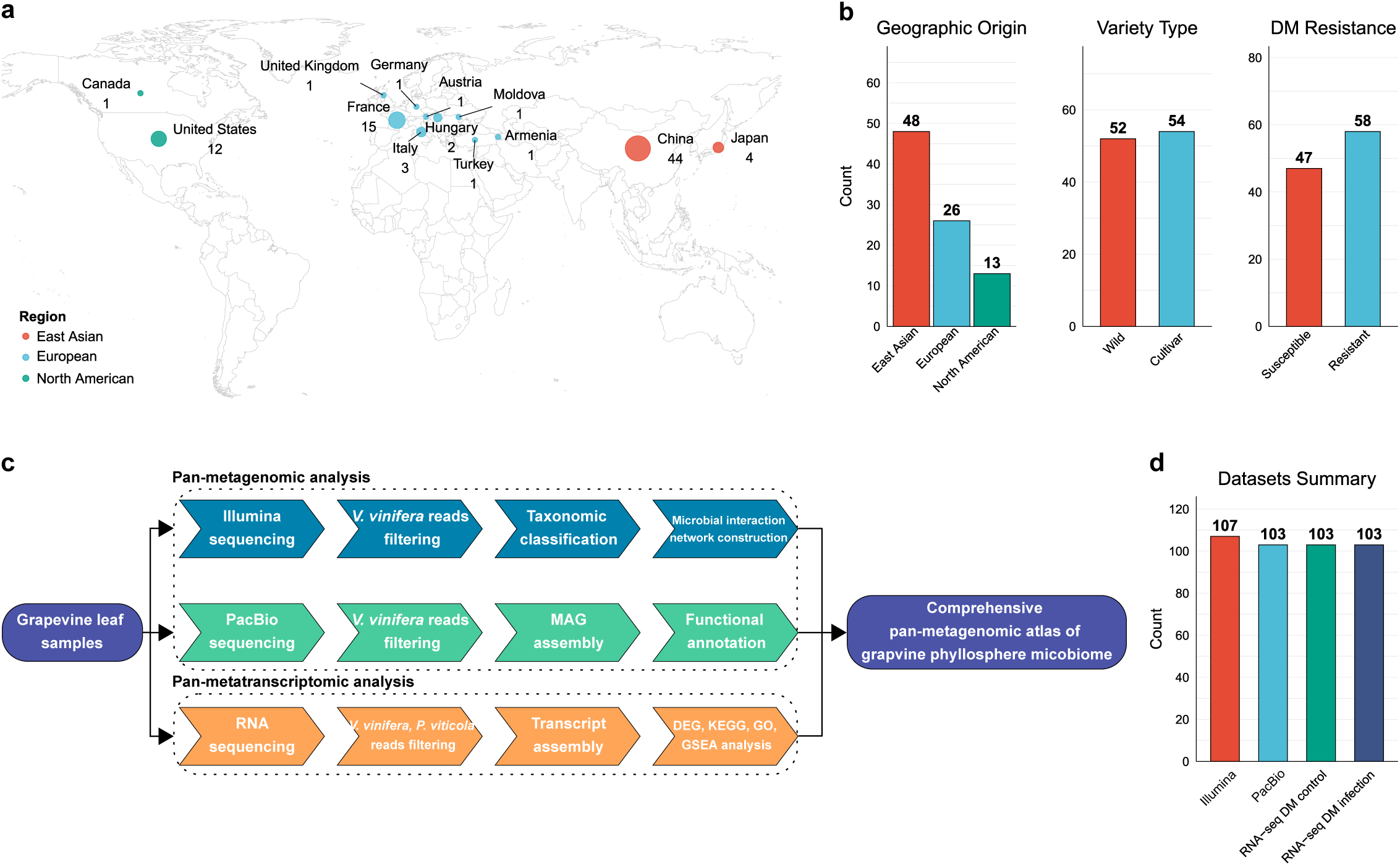
Research design for grapevine phyllosphere pan-metagenomics and pan-metatranscriptomics. **a**, Geographical origins of the sampled grapevine accessions. Numbers of samples were numerically labeled. **b**, Classification statistics of grapevine accessions by geographical origin, cultivar type, and DM resistance status. **c**, Pan-metagenomic and pan-metatranscriptomic analysis workflow. Colored blocks indicate sequencing methods: blue (Illumina), green (PacBio), orange (RNA-seq), and purple (shared components). **d**, Summary of generated datasets.

**Fig. 2.**
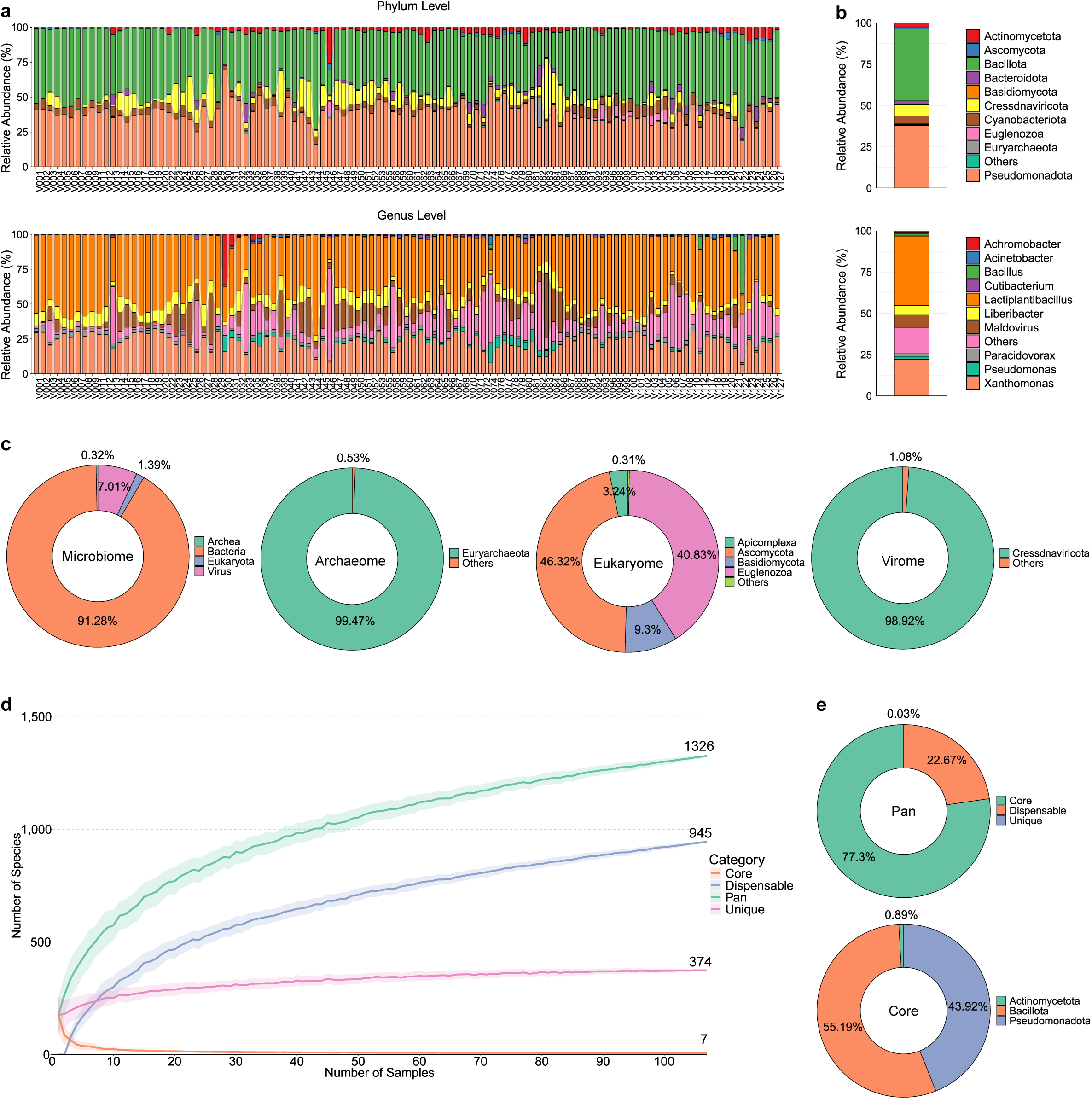
Pan-metagenomic landscape of the grapevine phyllospheric microbiome. **a**, Relative abundances of the 10 most abundant phyla (top) and genera (bottom) across the grapevine accessions. **b**, Mean relative abundance of the phyla and genera shown in (**a**). **c**, Compositional profiles of kingdoms from the grapevine accessions. **d**, Pan-metagenomic accumulation curve demonstrating pan, core, dispensable and unique genera across the grapevine accessions. **e**, Pan-metagenomic relative abundance at the genus level within the entire pan-metagenome (top) and the top 10 most abundant phyla within the core microbiome (bottom).

Although several previous studies have profiled grapevine microbiomes [11,14,17–24], mostly using 16S rDNA amplicon sequencing, a comprehensive pan-metagenomic analysis leveraging large-scale Illumina datasets had not been conducted prior to this study, which could provide critical insights into the structure and functional roles of grapevine microbiome. Through our pan-metagenomic analysis of 107 grapevine phyllosphere, we identified seven core genera (present in all accessions), 945 dispensable genera (present in some but not all accessions), and 374 unique genera (exclusive to a single accession) based on presence and absence patterns (**Fig. 2d and Table S3**). Strikingly, although the core microbiome accounted for only 0.6% of the detected genera, it dominated 77.3% of the relative abundance, with Bacillota (55.2%) and Pseudomonadota (44.0%) as the major phyla in the core microbiome (**Fig. 2e**). This result further underscores the central role of the core microbiome (mainly Bacillota and Pseudomonadota) in shaping the grapevine phyllosphere microbiome.

### Assembly of grapevine phyllospheric metagenome-assembled genomes (GPMAGs)

To assemble high-quality GPMAGs that provide further insights into the potential functions of the grapevine phyllosphere microbiome that potentially influence the host’s biology, we generated a total of 103 PacBio datasets with PacBio Sequel IIe system [27] (**Fig. 1d**) for the grapevine accessions used in Illumina-based metagenomic analysis above. The PacBio reads were mapped against the grapevine reference genome to completely exclude host-derived sequences, retaining unmapped reads representing microbial data (**Table S2**). These unmapped reads from all accessions were pooled together and then assembled into contigs by using various assemblers, such as Flye [30], hifiasm [31], hifiasm-meta [32], metaFlye [33], metaMDBG [34], and Verkko [35]. The contigs were subsequently binned using CONCOCT [36], Maxbin 2.0 [37], MetaBAT 2 [38], followed by dereplication with Galah (https://github.com/wwood/galah). This workflow generated a total of 19 non-redundant GPMAGs (**Fig. 3a and Table S4**) that met the medium-quality threshold based on standard MAG quality assessment criteria [34]. Among these, four were classified as complete, three as near-complete, five as high-quality, and seven as medium-quality MAGs (**Fig. 3b and Table S4**), highlighting the high quality of the GPMAGs. The median genome size of the GPMAGs were 4.5 Mb for the complete, 4.6 Mb for the near-complete, 4.7 Mb for the high-quality, 4.2 Mb for the medium-quality (**Fig. 3c and Table S4**), approximating typical bacterial genome sizes. Subsequently, GTDB-tk [39] was employed to identify the microbial profile of the GPMAGs. Among the classified GPMAGs, 63.2 % belonged to Pseudomonadota, 5.3% to Bacillota, and 21.1% to Actinomycetota (**Fig. 3d and Table S4**), which generally aligned with the Illumina-based microbial composition (**Fig. 2b**), except for Bacillota. To examine the pan-genomic profile of the GPMAGs among the different PacBio accessions, we further investigated the prevalence of the GPMAG with the different grapevine accessions measured by the PacBio read mapping rates against each GPMAGs. Across the grapevine accessions, *Mangrovibacter phragmitis* exhibited the highest prevalence, followed by *Roseateles* and *Sphingomonas aquatilis* (**Fig. 3e**). Interestingly, these GPMAGs all belonged to Pseudomonadota, further supporting its prevalence across grapevine accessions as indicated by the Illumina-based microbial composition (**Fig. 2b**). Moreover, all GPMAGs represent the highest MAG qualities (complete and near complete) also belong to Pseudomonadota, underscoring a strong association between MAG quality and prevalence.

**Fig. 3.**
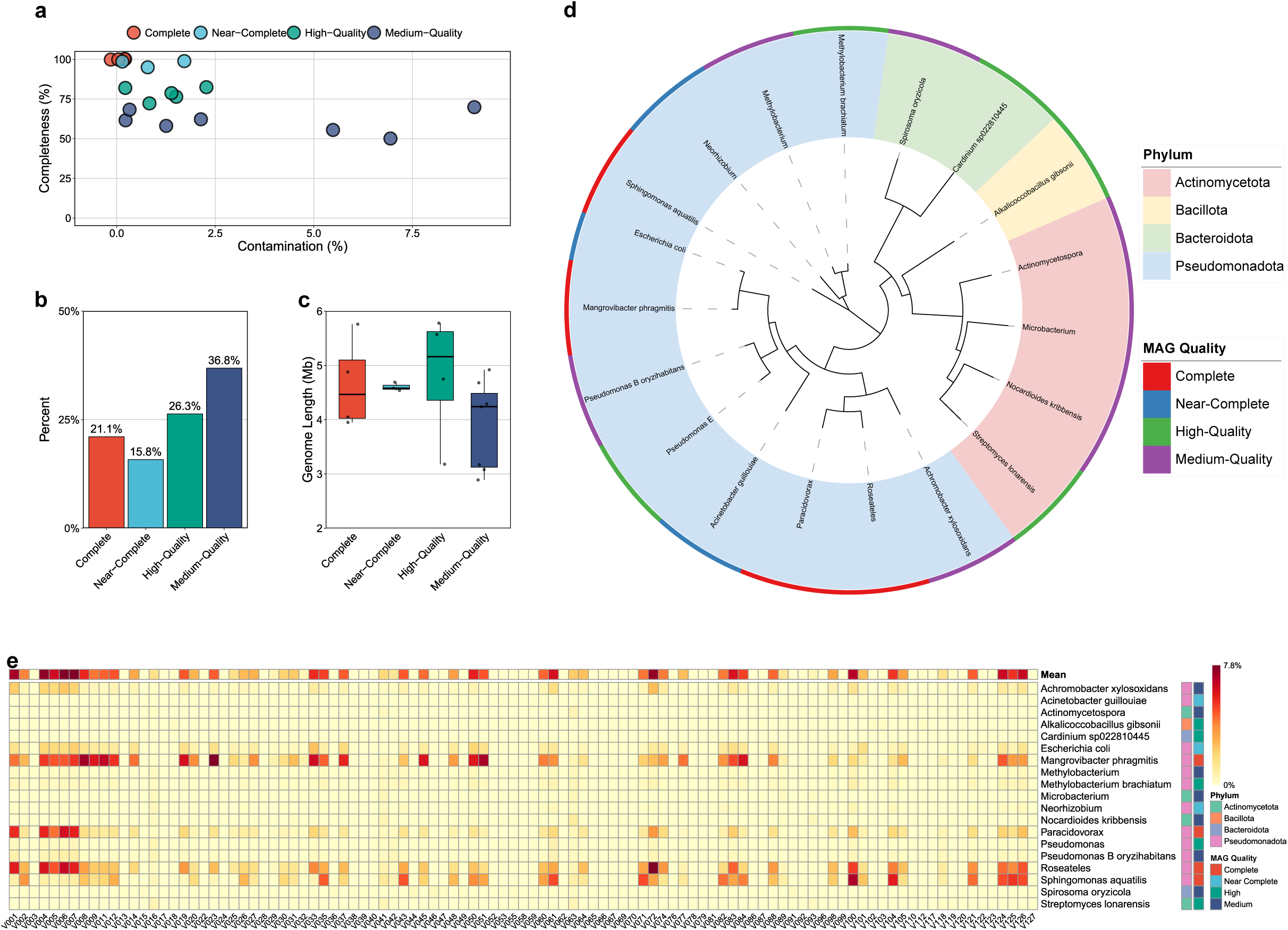
Assembly and genomic profile of the grapevine phyllospheric metagenome-assembled genomes (GPMAGs). **a**, Quality distribution of the GPMAGs in terms of MAG completeness and contamination. **b**, Composition of the GPMAG quality categories. **c**, Genomic size distribution of the GPMAGs. **d**, Phylogenetic tree of the classified GPMAGs, illustrating corresponding phyla and MAG qualities. **e**, Pan-genomic prevalence of the GPMAGs across the grapevine accessions. Heatmap depicts the relative abundance of the GPMAGs relative to each grapevine PacBio data accession.

Together, these results highlight high-quality and comprehensive assembly of PacBio-based GPMAGs representing valuable genomic resources for the grapevine phyllospheric microbiome investigation.

### Annotation of grapevine MAGs reveals genes encoding key biological processes

To delve deeply into functional roles of the GPMAGs in the grapevine microbiome, particularly those derived from unculturable microorganisms, we performed genomic annotation for the GPMAGs using Prokka [40] pipeline. This analysis identified a total of 82,884 genes, comprising 81,801 CDSs, 956 tRNA genes, 96 rRNA genes, 15 tmRNA genes, and 16 repeat regions (**Fig. 4a and Table S5**). Functional annotation of the CDSs was further conducted using eggNOG-mapper v2 [41], and annotated the detailed functions of the CDSs, which identified 67,012 CDSs with various biological process terms. In terms of biological function, proteins with unknown function (26.7%) represented the largest category among the CDSs, and the remaining was primarily associated with core biological functions, such as transcription, metabolism, and transport (**Fig. 4b**). Notably, the top 20 most abundant functional terms accounted for 93.9% of total terms, demonstrating a highly focused functional diversity in the GPMAGs that suggests specialized adaptation to the grape phyllosphere niche. Despite biosynthesis of secondary metabolites and microbial metabolism in diverse environments being the predominant biological pathway terms, the top 20 terms represented only 47.9% of total annotations (**Fig. 4c**), demonstrating a functional versatility in the GPMAGs. These results indicate both functional consistency and expansive metabolic potentials in the GPMAGs, likely facilitating phyllosphere colonization and adaptation in grapevines.

**Fig. 4.**
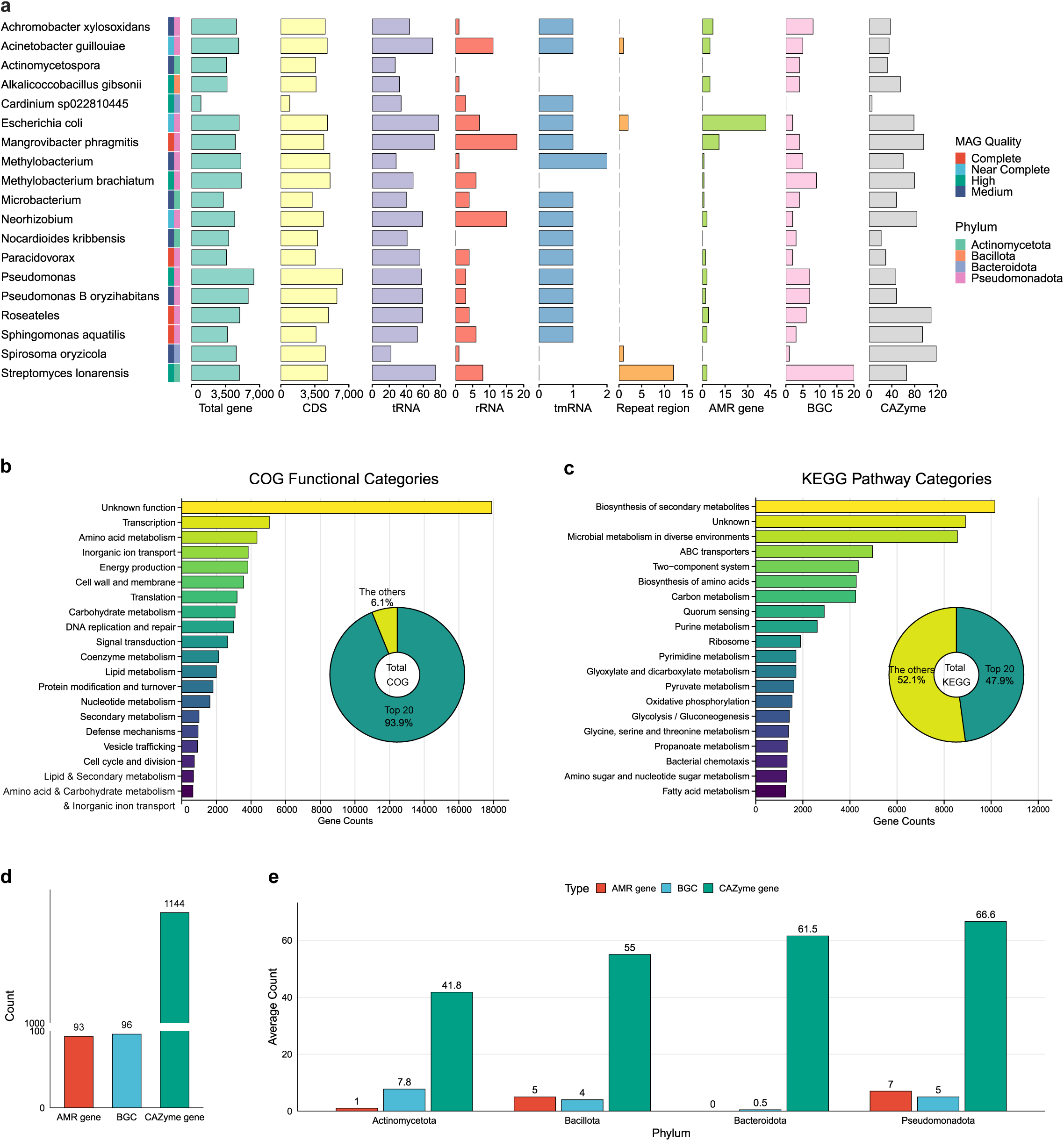
Functional annotation of the GPMAGs. **a**, Gene type distributions in the GPMAGs. **b-c**, Top 20 COG (**b**) and KEGG pathway (**c**) profiles of the GPMAGs. Donut plots displaying proportions of top 20 terms within total terms. **d**, Statistical summary of AMR, BGC, andCAZyme-associated genes. **e**, Phylum-level functional capacity displayed as mean AMR/BGC/CAZyme-associated gene counts per phylum.

Agriculture such as viticulture, heavily relies on pesticide application in disease management regime, but their misuse may have inadvertently contributed to the emergence of antimicrobial resistance (AMR) in the crop microbiome [42], posing significant threats to both food safety and public health. Notably, the annotated CDSs from GPMAGs frequently encode proteins associated with antibiotic resistance, such as ABC transporter (**Fig. 4c**), indicating the widespread presence of AMR genes within the GPMAGs. Specifically, we employed RGI [43] to identify 93 AMR genes spanning 16 distinct AMR families (**Fig. 4a, d, S2 and Table S6**). Notably, a significant proportion of the AMR genes exhibited multi-drug resistance through antibiotic efflux, a predominant resistance mechanism. The abundant presence of AMR genes in the grapevine microbiome implies that antibiotic resistance may have occurred due to prolonged pesticide application and misuse in viticulture.

Plant-associated bacterial endophytes protect their plant hosts from external pathogens by producing secondary metabolites (SMs), including antibiotics and toxins [44–46], establishing them as valuable resources for environmentally friendly and pathogen-effective biocontrol agents. Biosynthetic gene clusters (BGCs) responsible for bacterial SM production typically comprise dozens of related genes distributed across extensive genomic regions, necessitating high-quality MAGs with exceptional completeness and continuity for comprehensive exploration. To leverage our high-quality GPMAGs, we employed antiSMASH [47] to identify a total of 96 BGCs from 26 distinct BGC types in the grapevine phyllospheric microbiome (**Fig. 4a, d, S3 and Table S7**). Among these, terpene, RIPP-like, NRPS clusters were the most abundant, commonly associated with antimicrobial activity critical for host defense [48–50]. Other predominant BGCs, including Ni-siderophore, betalactone, and T3PKS highlight the microbiome’s ample potentials in SM productions. These candidate BGCs provide valuable resources for discovering novel natural products and elucidating their roles in grapevine biology and medicinal applications.

Several studies have demonstrated that microorganism-derived polysaccharides play crucial roles in plant bioprocesses, including immune responses, growth facilitation, stress adaptation, photosynthesis, and plant-bacteria interactions [51–53]. To explore the potential of polysaccharides production in the grapevine microbiome, we employed dbCAN [54] to annotate CAZyme genes from the GPMAGs. The analysis identified a total of 1,144 CAZyme genes spanning 143 unique enzyme families, reflecting the grape phyllospheric microbiome’s substantial capacity for polysaccharides metabolism (**Fig. 4a, d, S4 and Table S8**). Among these genes, glycosyltransferases (GTs) and glycoside hydrolases (GHs) were particularly abundant, with GT2, GT4, GT51, and GH13 emerging as the most prevalent, suggesting a strong focus on polysaccharide synthesis and modification. Additionally, the presence of diverse carbohydrate-binding modules (CBMs) and auxiliary activities (AAs) highlights the microbiome’s versatility and its potential involvement in plant-microbe interactions. This extensive CAZyme repertoire indicates that the grape phyllospheric microbiome may influence various host physiological processes, including disease resistance and growth, through polysaccharide synthesis and other host-interactive mechanisms.

To evaluate the functional potentials of each phylum in terms of antibiotic resistance, secondary metabolite biosynthesis, and carbohydrate metabolism that differentially influence the host’s biology, we quantified the mean numbers of AMR genes, BGCs, and CAZyme genes of each phylum in the GPMAGs. As a result, Pseudomonadota contained the highest abundance of AMR genes (**Fig. 4e**), indicating particularly strong antibiotic resistance potential. Actinomycetota showed the greatest capacity for secondary metabolite biosynthesis, harboring the most BGCs. While all phyla maintained comparable CAZyme gene numbers, Pseudomonadota demonstrated particularly extensive carbohydrate metabolic potential. These results reveal phylum-specific functional potentials that differentially involve the host’s biological processes.

In conclusion, these findings provide valuable insights into the functional potential of the grapevine phyllospheric microbiome, emphasizing its involvement in shaping the host health and resilience through microbial-host interactions.

### Downy mildew resistance relates to complexity of microbial interaction network with presence of key genera

To investigate the role of grapevine microbiome during *P. viticola* infection, we profiled the microbiome composition of the grapevine accessions based on their DM disease phenotypes (susceptible or resistant). Unexpectedly, the microbiome compositions of the susceptible and resistant microbiomes were nearly identical (**Fig. S5a**). Additionally, β-diversity analysis further revealed no overall compositional difference between the two groups (**Fig. S5b**), suggesting that microbiome composition unlikely accounts for the differences in DM resistance. Therefore, we hypothesized that interactions within the microbial community might contribute to the observed resistance. To test this hypothesis, we analyzed the complexity of the microbial interaction networks in the susceptible and resistant grape microbiome using Sparse Inverse Covariance estimation for Ecological Association and Statistical Inference (SpiecEasi) [55] analysis. SpiecEasi uses sparse inverse covariance estimation to identify true associations between microorganisms based on correlations derived from the covariance of their relative abundances. Positive correlations indicate cooperative interactions, while negative correlations represent antagonistic interactions. Remarkably, the microbiome associated with resistant plants exhibited a significantly more complex network than the susceptible ones (**Fig. 5a**), with a higher frequency of negative interactions per genus (**Fig. S6**), indicating the involvement of microbial network complexity in modulating DM-resistance, especially the negative interactions play crucial role for the complexity.

**Fig. 5.**
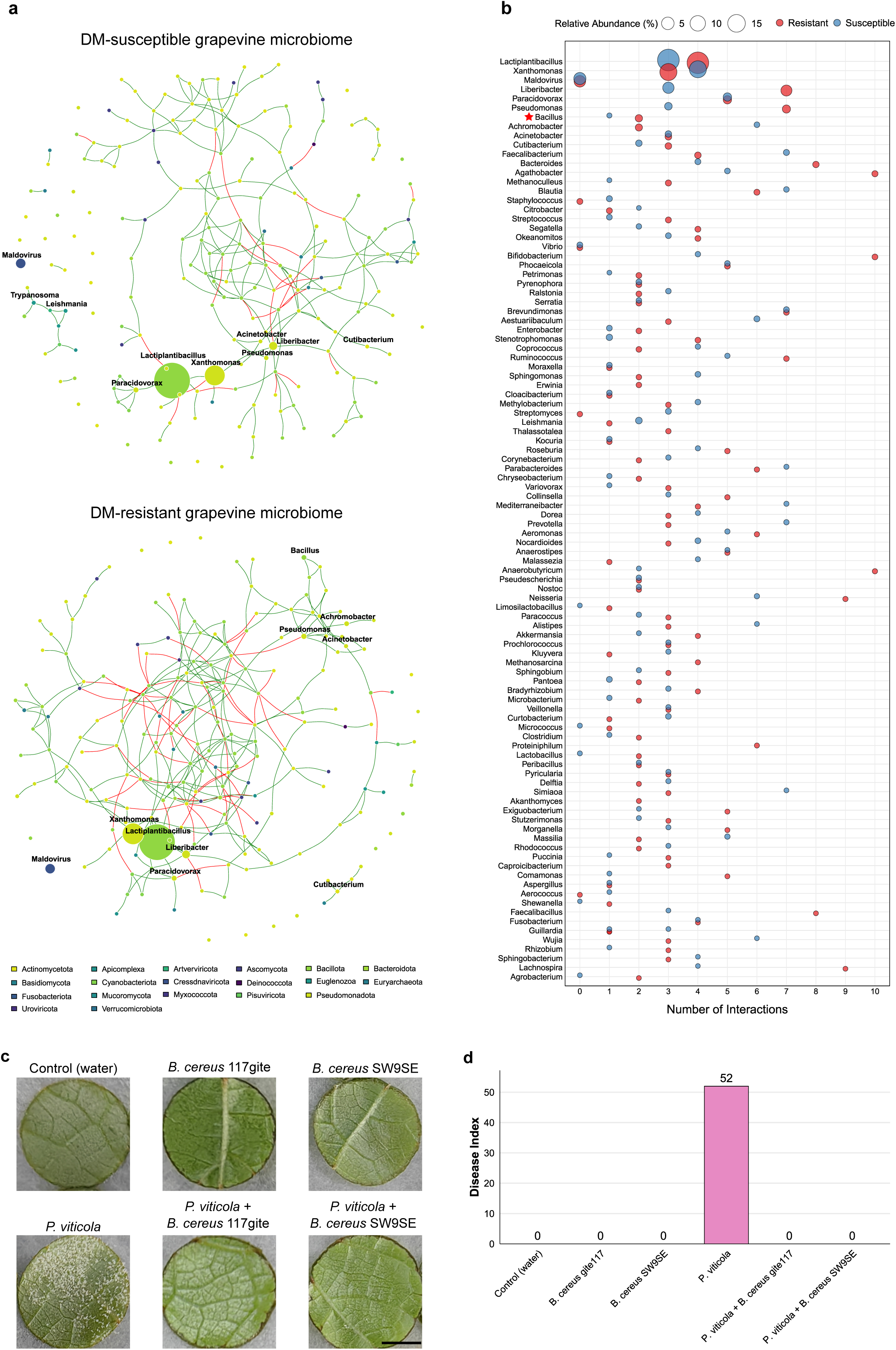
Downy-mildew resistance shapes complex microbial interaction networks with key genera. **a**, Microbial co-occurrence networks based on the DM resistance types. The networks displaying the top 200 most abundant genera within each type. Node sizes indicate genera’ relative abundances, with colors denoting different phyla as shown in bottom panel legend. Green edges represent positive correlations, while red edges indicate negative correlations between nodes. The top 10 abundant genera within each microbiome were labeled above corresponding nodes. **b**, Profiles of top 100 abundant key microorganisms in the DM-resistant grapevine microbiome, illustrating relative abundances and numbers of microbial interactions. Colored bubbles represent the profiles in the corresponding microbiome. The top 100 genera of the resistant microbiome which not present in the top 200 genera in the susceptible microbiome were not shown in the plot. Red star indicates a genus that experimentally isolated and validated in this study. **c**, Biocontrol assay of two *B. cereus* strains against *P. viticola* infection. The top and bottom rows show mono- and co-inoculation, respectively. Black bar indicates 5 mm. **d**, Bar plot of the disease index shown in (**c**), where higher values indicate more severe infection.

Notably, the microbiome associated with resistant plants was highly compartmentalized and centralized around a few genera (e.g., *Lactiplantibacillus*, *Liberibacter, Paracidovorax*, and *Xanthomonas*), with relatively much higher abundance (**Fig. 5a**), suggesting the presence of key genera with pivotal roles in network dynamics and resistance modulation. To identify these key genera, we first profiled the top 100 most abundant genera in the resistant microbiome based on relative abundance and microbial interaction metrics. These profiles were then systematically compared with those from the susceptible microbiome to evaluate their respective contributions to the network complexity. Among them, *Liberibacter* and *Pseudomonas* emerged as the most influential genera, with significantly increased interaction metrics compared to those of the susceptible microbiome (**Fig. 5b**). Intriguingly, all these key genera belong to the core microbiome (**Table S3**), highlighting the importance of the core microbiome in modulating the complexity. Additionally, genera such as *Bacillus* exhibit increased abundance and interaction metrics compared to the susceptible microbiome, indicating contribution to the complexity via abundance and microbial interaction. Despite relatively low abundances in the resistant microbiome, genera such as Bacteroides, Agathobacter, Bifidobacterium, Anaerobutyricum, and Neisseria exhibit the highest interactions (8-10 interactions/genus) in the resistant microbiome, demonstrating the important role of low abundant genera in forming the complexity. To evaluate phylum-level contributions to the increased complexity, we compared the proportions of microbial interaction between the resistant and susceptible microbiomes for each phylum. As a result, Bacillota was identified as the primary contributors exhibiting the most significantly increased proportion in the resistant microbiome (33.5%) compared to that of the susceptible one (23.9%), while other phyla show minimal fluctuation in the change (**Fig. S7**). Furthermore, we analyzed Spearman correlations between Bacillota and other major phyla in the resistant microbiomes based on their abundances, identifying those with the strongest interactions with this key phylum. The correlation analysis revealed that Euglenozoa, one of the major eukaryotes in the grapevine microbiome (**Fig. 2c**) and Cyanobacteriota exhibits the strongest positive interaction with Bacillota, while the other phyla display no significant positive interactive dynamics (**Fig. S8**), indicating that Bacillota positively interacts with these phyla in the resistant microbiome. These findings suggest that not all genera contribute equally to the modulation of the network complexity; rather, specific key genera play a primary role along with specific interactions.

To validate the effectiveness of key genera profiling and role of key genera during *P. viticola* infection, we screened microorganisms isolated from the grapevine leaves. Among the key genera, we successfully identified two *Bacillus* strains which belong to Bacillota, *B. cereus* 117 gite and *B. cereus* SW9SE. Given our observation that *Bacillus* is a key genus belong to Bacillota with increased proportion and microbial interactions in the microbiome associated with resistant plants (**Fig. 5b and S5a**), and considering a prior report on *B. cereus* efficacy against grape ripe rot disease [56], we hypothesized and tested the biocontrol potential of *B. cereus* against *P. viticola* infection. To this end, we inoculated grapevine leaf discs with varying concentrations of these strains followed by *P. viticola* infection. Strikingly, both strains exhibited apparent DM inhibition at a low concentration (e.g., OD=0.02) (**Fig. 5c and d**), demonstrating the potent biocontrol activity of the strains against *P. viticola* infection. These findings underscore the effectiveness of our key genera profiling, and paving a way for *B. cereus* as a novel biocontrol resource for managing DM infection in viticulture.

Collectively, our pan-metagenomic analysis and experiments highlight the association of the microbial network complexity with DM-resistance, driven by intricate interactions involving specific key genera with specific interactions.

### Downy mildew triggers eukaryotic primary and secondary metabolism of the grapevine microbiome

Given that network complexity is primarily driven by Bacillota, which exhibits the strong positive abundance correlation with one of the dominating Eukaryotes-Euglenozoa, we hypothesize that key bacterial phyla involve eukaryotic transcriptomes in the grapevine microbiome, potentially underlying defensive responses against *P. viticola* infection. To examine this, we conducted pan-metatranscriptomic sequencing and analysis to investigate the transcriptional regulation of the grapevine phyllospheric microbiome in response to the infection. In brief, we collected samples from both infected (one day post-inoculation) and uninfected leaves from 103 of the grapevine accessions and subjected them to RNA-seq [27] (**Fig. 1d**). The RNA-seq reads were mapped to the grapevine and *P. viticola* reference genomes (NCBI accession: GCA_001695595.3), obtaining the unmapped reads that represent eukaryotic organisms other than grapevine and *P. viticola* (**Table S2**). The unmapped reads were then *de novo* assembled using Trinity [57], resulting in 192,675 eukaryotic transcripts. Differentially expressed transcript (DET) analysis identified 40,546 significantly up-regulated and 5,081 down-regulated transcripts (**Fig. 6a**), indicating a considerable overall upregulation of the microbial transcripts upon the infection. Moreover, KEGG and GO enrichment revealed that the up-regulated genes were primarily involved in primary and secondary metabolisms (**Fig. 6b and c**), with secondary metabolism being particularly prominent in the microbiome’s response to the infection. Consistently, Gene Set Enrichment Analysis (GSEA) further confirmed significant enrichment of secondary metabolism-related gene sets among the upregulated transcripts (**Fig. 6d**). Notably, these upregulated transcripts encoded the key proteins of BGCs, including ketoacyl reductase (KR) domains and flavin mononucleotide-dependent dehydrogenases, both of which are integral to the biosynthesis of polyketides and antibiotics [58,59]. Together, these findings demonstrate that primary and secondary metabolisms in eukaryotic microorganisms are significantly induced during *P. viticola* infection, suggesting a potential role for the grapevine phyllospheric microbiome in defending against pathogen invasion through the competitive exclusion and SM productions that associated with key bacterial genera.

**Fig. 6.**
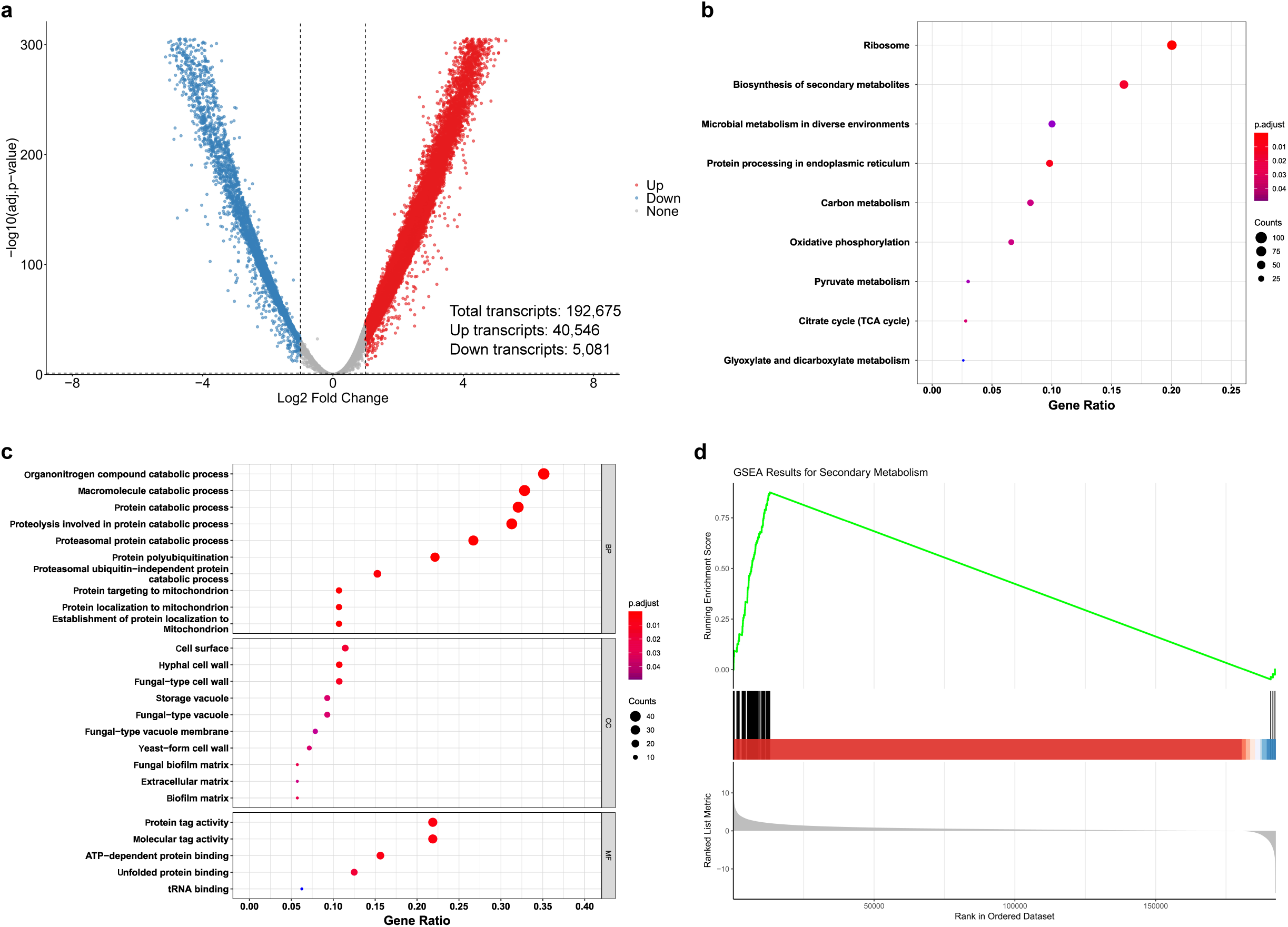
Eukaryotic transcriptome profile of the grapevine phyllospheric microbiome in response to the oomycete Plasmopara viticola infection. **a**, Volcano plot of the grape microbiome upon infection by P. viticola. The dashed lines indicate thresholds for statistical significance (log2 fold change > 1, adjusted p < 0.05). **b-c**, KEGG pathway (**b**) and GO enrichment analysis (**c**) of the up-regulated DETs. GO enrichment shows the top 10 terms in each category. **d**, GSEA of the secondary metabolism-related transcripts. The graph illustrates the enrichment score (green line) for the transcript set associated with secondary metabolism across the ranked transcript list (bottom) by log2 fold change values. The black bars in the middle represent the positions of transcripts from the set in the ranked list with significances indicated by red and blue.

## Discussion

Microbiome have profound impact on physiology and immunity of host organisms ranging from human, animals and plants. Mounting evidence underscores the critical roles of the microbiome in the health, yield, and quality [4,11–13] of grapevine which is one of the most economically important fruit crops cultivated worldwide for production of fresh fruits, wine, juice and raisin etc. Despite a huge body of knowledge gained in microbiome research, pan-microbiome diversity across host genera is poorly understood, lacking a large-scale pan-metagenomic approach, especially from long-read sequencing. In this study, combining short, long-read sequencing and RNA-seq, we conducted a pan-metagenomic and pan-metatranscriptomic study of 107 accessions from 34 *Vitis* species to understand the structure, diversity and functional roles of pan-microbiome in grapevine phyllosphere, a key battleground for host-microbe interactions. We identified the core microbiome dominated by Bacillota and Pseudomonadota. Leveraging PacBio sequencing of grapevine phyllosphere, we successfully assembled 19 high-quality GPMAGs including four near-complete MAGs, with detailed functional annotations, representing the first assembly of MAGs derived from plant PacBio datasets. Furthermore, pan-metagenomic and pan-metatranscriptomic analyses elucidated the underlying defense mechanisms of grape microbiome against *P. viticola* infection. Our findings demonstrate the profound influence of the microbiome on grapevine biology, particularly involvement in the resistance of *P. viticola* infection, and provide a foundation and valuable genomic and microbial resources for developing microbiome-based biocontrol strategies in viticulture.

Structural analysis of the grapevine phyllosphere microbiome revealed the core microbiome dominated by specific phyla, primarily Bacillota and Pseudomonadota, which constitute the dominant proportion of the microbial community (**Fig. 2**). This composition aligns well with previous studies on grapevine phyllosphere microbiota across different cultivars and geographic regions [4,15,60,61], highlighting the prevalence and pivotal roles of the core microbiome. However, the mechanisms underlying core microbiome recruitment in the grapevine phyllosphere remain largely unknown. In plants, it is well established that metabolic secretions, particularly SMs such as flavonoids, salicylic acid, and strigolactones, shape microbiome composition through plant-microorganism interactions [62–65]. In turn, the modulated microbiome significantly influences host biological processes via plant-microbe interaction, including growth, disease resistance, and stress responses [62–65]. As a major plant part interacting with microbial communities, common SMs might accumulate in grapevine leaves of diverse genetic background, thereby helping establish the core microbiome. This core microbiome plays critical roles in growth, disease resistance, and stress responses, underscoring the importance of grapevine-microbe interactions in maintaining normal plant development.

Compared to amplicon sequencing analysis, MAGs provide novel and pivotal insights into plant microbiome. Our GPMAGs demonstrate the high quality and prevalence, serving as ideal genomic resources with detailed annotations such as AMR, BGC, and CAZyme information for further applications. To our knowledge, this is the first study to generate MAGs from PacBio reads of a large collection of plant accessions. However, the rare presence of dominant phyla such as Bacillota in the GPMAGs (**Fig. 3d and Table S4**) highlights the limitations of long-read technology in the number of microbial genomes recovered from microbiome, primarily due to higher cost of sequencing and thus lower data coverage compared to Illumina data, and limitation of bioinformatic methods [66,67]. Decreasing sequencing cost and further optimization and advancement of these methods will improve PacBio-based MAG assemblies in the future. Together, this study establishes a novel and robust method for assembling MAGs from plants, leveraging the contiguity and accuracy of PacBio reads to accelerate research on plant microbiome.

It is widely recognized that through interactions microorganisms, plants modulate their immunity to combat pathogen invasion [68,69]. This mechanism primarily relies on microbiome-derived SM production and competitive exclusion to protect host plants from various pathogens. Consistent with this concept, our findings demonstrate the pivotal roles of bacteriome and eukaryome, of which bacteriome re-orchestrates microbial community, especially facilitates interactions with microorganisms to involve their SM production and competitive exclusion in response to *P. viticola* infection (**Fig. 7**). Intriguingly, we observed a significant increase in network complexity, driven by the contributions of key genera from bacteriome (**Fig. 5 and Fig. S6, 7**), which involves the defense mechanism against *P. viticola* infection in grapevines through interacting with eukaryome. In olive, *Pseudomonas simiae* PICF7, a biological control agent for Verticillium wilt, was reported to significantly increase microbial network complexity upon the infection, resulting in a more stable network that plays a crucial role in defense [70]. Thus, it is likely that grapevines employ a similar strategy to modulate their microbial network, optimizing SM production and competitive exclusion capacities of eukaryome to effectively combat *P. viticola* infection. Additionally, the profiling of key genera in the DM-resistant microbiome provides a valuable resource for exploring biocontrol microorganisms against *P. viticola* infection exemplified by the experimental validation of two *B. cereus* strains, laying the foundation for environmentally friendly and sustainable biocontrol strategies in viticulture.

**Fig. 7.**
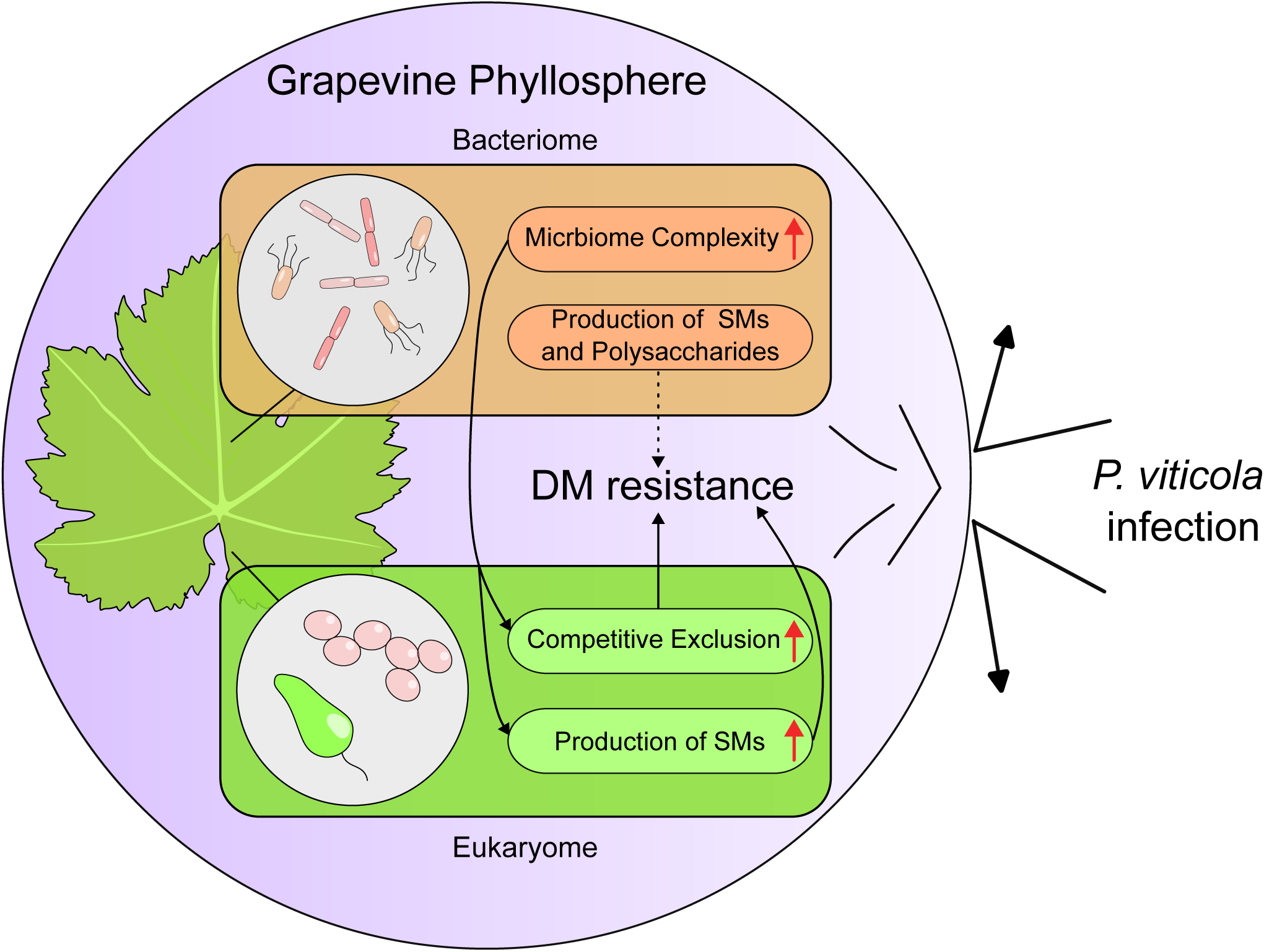
Schematic diagram depicting grapevine microbiome’s potential deployment of bacteriome and eukaryome to combat *P. viticola* infection. This pan-metagenomic and pan-metatranscriptomic study demonstrates that the grapevine bacteriome orchestrates microbiome assembly, with key taxa (e.g., Bacillota) interacting selectively with eukaryotes (e.g., Euglenozoa) to involve eukaryotic competitive exclusion and secondary metabolites (SMs). The bacteriome may further utilize SMs and polysaccharides for enhanced DM resistance (dashed line, speculative).

## Materials and methods

### Collection of grapevine leaves for metagenome sequencing

The leaf samples from 107 grapevine accessions used in this study were collected from vineyards at the Zhengzhou Fruit Research Institute, Chinese Academy of Sciences (Zhengzhou, Henan, China), and the Institute of Advanced Agricultural Sciences, Peking University (Weifang, Shandong, China). All grapevines were maintained under standard management practices, including watering, fertilization, pruning, and disease control.

### Downy mildew infection assay

DM infection was performed as described previously [71]. Each grapevine leaf disc (8 mm in diameter) was inoculated with 20 μL of *P. viticola* suspension (1×10⁵ sporangia/mL) on the abaxial surface and incubated at 18 ± 2°C with 80 ± 10% relative humidity in a growth chamber under a 12 h light (60 μmol·m⁻²·s⁻¹)/12 h dark cycle. Leaf discs were collected 24 hours post-inoculation for transcriptome analysis. At least three biological replicates were conducted, with each replicate comprising five leaf discs from different grapevines.

### DNA and RNA extraction

Genomic DNA for Illumina was extracted from the grapevine leaves using DNeasy Plant Mini Kit (QIAGEN, Hilden, Germany) according to the manufacturer’s instructions. The DNA integrity was assessed with an Agilent 4200 Bioanalyzer (Agilent Technologies, Santa Clara, USA). High-molecular-weight (HMW) DNA for PacBio sequencing was prepared from the fresh grapevine leaves using the cetyltrimethylammonium bromide (CTAB) method [72]. The DNA quality was verified with 1% agarose gel electrophoresis, and purity was assessed using a NanoPhotometer® spectrophotometer (IMPLEN, Westlake Village, USA). The DNA concentration was measured with a Qubit® 2.0 Fluorometer (Life Technologies, Carlsbad, USA). Total RNA was extracted using TRIzol reagent (Thermo Fisher Scientific, Waltham, USA) following the manufacturer’s instructions. The RNA concentration was measured with a Nanodrop 2000 spectrophotometer (Thermo Fisher Scientific, Waltham, USA), and the purity was evaluated using a NanoPhotometer® (IMPLEN, Westlake Village, USA). The RNA concentration and integrity were further analyzed with an Agilent 2100 Bioanalyzer using the RNA Nano 6000 assay kit (Agilent Technologies, Santa Clara, USA).

### Metagenome sequencing

For Illumina library preparation, 15 µg of the genomic DNA was sheared using g-Tubes (Covaris, Woburn, USA) and concentrated with AMPure PB magnetic beads (Pacific Biosciences, Menlo Park, USA). The library construction was performed with 1 µg of the DNA template following TruSeq DNA Sample Preparation Guide (Illumina, San Diego, USA). Paired-end sequencing was conducted on a HiSeq X Ten platform (Biomarker Technologies, Qingdao, China), generating 150 bp Illumina reads. For PacBio sequencing, 15 µg of the purified HMW genomic DNA was used to construct a standard SMRTbell library with SMRT Express Template Prep Kit 2.0 (Pacific Biosciences, Menlo Park, USA). Circular consensus sequencing (CCS) producing PacBio reads was performed on a PacBio Sequel IIe system at Biomarker Technologies (Qingdao, China). To prepare RNA sequencing libraries, 1-3 µg of the total RNA was used as input for library construction with VAHTS Universal V6 RNA-seq Library Prep Kit for Illumina® (Vazyme, Nanjing, China) according to the manufacturer’s instructions. The RNA concentration was initially quantified with the Qubit® RNA Assay Kit on a Qubit® 3.0 (Thermo Fisher Scientific, Waltham, USA) and diluted to 1 ng/µl. Insert size was measured with an Agilent Bioanalyzer 2100 (Agilent Technologies, Santa Clara, USA), and final library concentration (>10 nM) was verified with a Bio-RAD CFX 96 quantitative PCR system (Bio-Rad, Hercules, USA). Paired-end sequencing was then performed on a HiSeq X Ten platform (Biomarker Technologies, Qingdao, China), generating 150 bp RNA-seq reads.

### Quality control and removal of host reads

To ensure the quality of the short reads, quality control was performed on the Illumina and RNA-seq reads using fastp [73] (v0.23.4) with default parameters, resulting in the quality-controlled reads. To completely filter out reads from the grapevine hosts, the Illumina, PacBio, and RNA-seq reads were mapped to the complete reference genome of *Vitis vinifera* [27] (VHP-T2T.hap1, NCBI accession: GCA_046255155.1) using Minimap2 [74] (v2.28) with parameters (Illumina, RNA-seq: -ax sr, PacBio: -ax map-pb). The mapped results were further filtered using SAMtools [75] (v1.21) with parameters (Illumina, RNA-seq: view –u -f 12 -F 256, PacBio: view -f 4 -F 256) to retain microbial reads. Additionally, the RNA-seq reads were further filtered by mapping to *P. viticola* reference genome INRA_Pvit_2 (NCBI accession: GCA_001695595.3) with Minimap2 and SAMtools to exclude *P. viticola* reads.

### Taxonomic classification

To profile the taxonomic composition of the grapevine microbiome, the filtered Illumina reads were taxonomically classified using Kraken2 [28] (v2.1.3) and k2_pluspf_20240904 database with parameters (--paired --confidence 0.1 --minimum-hit-groups 2), yielding the compositional statistics of the grapevine microbiome.

### Pan-metagenomic and diversity analysis

The identification of the core, dispensable, and unique microbiome was conducted with R using the microbial composition statistics. To assess microbiome diversity, PCoA analysis was performed with vegan [76] (v2.6-8) package in R. All statistical analysis and visualization were performed with R.

### Assembly of the GPMAGs

To generate the GPMAGs, the filtered PacBio reads were assembled using Flye [30] (v2.9.5), Hifiasm [31] (v0.19.9), Hifiasm-meta [32] (vhamtv0.3.2), metaFlye [33] (v2.9.5), MetaMDBG [34] (v1.0), and Verkko [35] (v2.2.1) with default parameters to produce initial contig assemblies. The resulting contigs were then binned with CONCOCT [36] (v1.1.0), MaxBin2 [37] (v2.2.7), and MetaBAT2 [38] (v 2.17) with default parameters. The bins were quality-checked using CheckM2 [77] (v1.0.2), and those meeting the medium-quality criterion [34] were dereplicated to generate non-redundant GPMAGs using Galah (v0.4.2) (https://github.com/wwood/galah) with parameters (--ani 95 --precluster-ani 90 --min-aligned-fraction 30 --min-completeness 50 --max-contamination 10).

### Taxonomic and phylogenetic analysis of the GPMAGs

To reveal the taxonomic composition of the GPMAGs, GTDB-Tk [39] (v2.4.0) was employed with the R220 database and default parameters, generating the taxonomic composition and a multiple sequence alignment (MSA) file. The MSA file was further subjected to the construction of phylogenetic tree using IQ-TREE2 [78] (v2.1.4_beta) with default parameters, and visualized by iTOL [79] (https://itol.embl.de/).

### Functional annotation of the GPMAGs

To analyze different gene types from the GPMAGs, Prokka [40] (v1.14.6) was used for gene type prediction with default parameters. The predicted CDSs were further applied for functional annotation using eggNOG-mapper [80] (2.1.12) with parameters (-m diamond - -evalue 1e-5 --query_cover 50 --subject_cover 50). To annotate AMR genes, BGCs, and CAZyme genes of the GPMAGs, RGI [43] (v6.0.3) (default parameters), antiSMASH [47] (v7.1.0) (parameters: --genefinding-tool prodigal --taxon bacteria –fullhmmer --cb- knownclusters --cb-subclusters --cc-mibig --asf --rre --pfam2go), and dbCAN [54] (v4.1.4) (default parameters) were employed, respectively.

### Construction of microbial interaction network

Microbial interaction network was constructed using the microbial composition statistics and SpiecEasi [55] (v1.1.3) package in R with parameters (lambda.min.ratio=0.5, nlambda=30, rep.num=100).

### Isolation of grapevine phyllosphere microorganisms

To isolate microorganisms from grapevine leaves, we collected 10 leaves from each of the following cultivars: *V. vinifera* cv. Chardonnay, *V. vinifera* cv. Muscat Hamburg, *V. vinifera* x *V. labrusca* cv. Kyoho, and *V. vinifera* x *V. labrusca* cv. Vidal Blanc at the vineyards. After rinsing with sterile water, the leaves were homogenized in 1x PBS buffer using an electric mixer to obtain a microbial suspension. 2 ml of the suspension were inoculated onto Nutrient Agar medium (Coolaber Technology Co., Ltd., Beijing, China) and incubated at 30°C for 24 hours. Single colonies were isolated and further purified by streaking on Nutrient Agar medium. Species identification was performed by 16S rRNA sequencing (Tsingke Biotechnology Co., Ltd., Beijing, China).

### Experimental validation of biocontrol activity

To validate the efficacy of isolated microorganisms against *P. viticola* infection, the *B. cereus* strains were cultured in Nutrient Broth medium (Coolaber Technology Co., Ltd., Beijing, China) at 30°C and 220 rpm for 12 hours. The cultures were diluted to OD of 0.02 at 600 nm using a Nanodrop 2000 spectrophotometer (Thermo Fisher Scientific, Waltham, USA). The diluted suspensions were sprayed onto 1 cm grape leaf discs and incubated at 30°C and 220 rpm for 24 hours. Subsequently, *P. viticola* was co-inoculated onto the discs using the method described above and incubated at 19°C for 6 days. Disease severity was assessed 6 days post-inoculation using the following grading scale: Grade 0, no lesions (0% of leaf disc area affected); Grade 1, lesions covering less than 10% of the leaf disc area; Grade 2, lesions covering 10-30% of the leaf disc area; Grade 3, lesions covering 31-60% of the leaf disc area; Grade 4, lesions covering 61-80% of the leaf disc area; and Grade 5, lesions covering more than 80% of the leaf disc area. The disease index (DI) was calculated as follows:

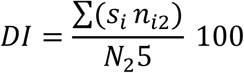

where *s_i_* represents the disease grade (ranging from 0 to 5), *n_i2_* is the number of leaf discs with disease grade *s_i_*, and *N*_2_ is the total number of leaf discs assessed. The DI value was rounded to one decimal place.

### Transcriptomic analysis of *P. viticola* infection

Transcriptomic profiling in response to *P. viticola* infection was conducted using the filtered RNA-seq reads. First, *De novo* assembly of transcripts was performed with Trinity [57] (v2.15.2) using default parameters. The generated transcripts were dereplicated using VSEARCH [81] (2.29.0) with parameters (--id 0.95), selecting the longest transcripts as the representatives of duplicates. Transcript expression was quantified using Salmon [82] (v1.10.3) with default parameters and the filtered RNA-seq reads. Differential expression analysis was performed with the DESeq2 [83] (v1.45) package in R. Subsequently, the significantly up-regulated genes were further annotated with eggNOG-mapper, followed by KEGG, GO, and GSEA enrichment analyses using AnnotationForge (v1.47.1), AnnotationDbi (v1.67.0), ClusterProfiler [84] (v4.13.3) and enrichplot (v1.25.2) packages in R with default parameters.

## Acknowledgments

We would like to thank the Bioinformatics Platform at Peking University Institute of Advanced Agricultural Sciences for providing the high-performance computing resources.

## Author contributions

LG and WXY conceived and supervised the study. JYJ, XNZ, JJM, WZ, LLG, HSS, CHL and LLZ prepared the plant samples and performed the experiments. JYJ and XFW conducted the bioinformatics analysis. JYJ, LG and WXY wrote and revised the manuscript. All authors read and approved the final manuscript.

---

**Fig. S1.**
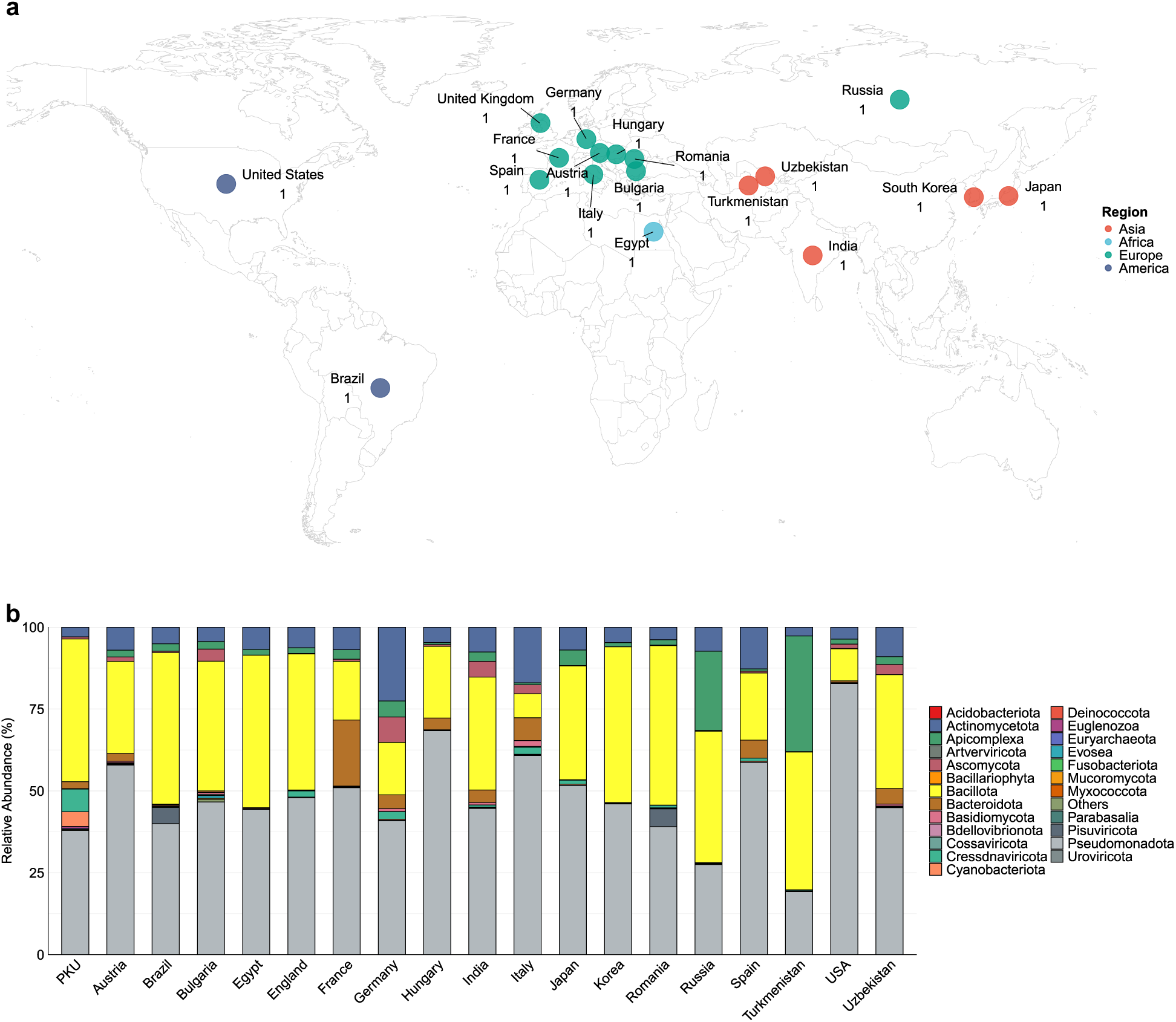
Worldwide comparison of grapevine phyllosphere microbiome compositions. **a**, Geographical distribution of Vitis accessions used for comparative analysis. One representative Vitis accession was selected per country. **b**, Top 10 phylum level composition of grapevine phyllosphere microbiome across worldwide Vitis accessions. PKU represents mean phylum composition of this study.

**Fig. S2.**
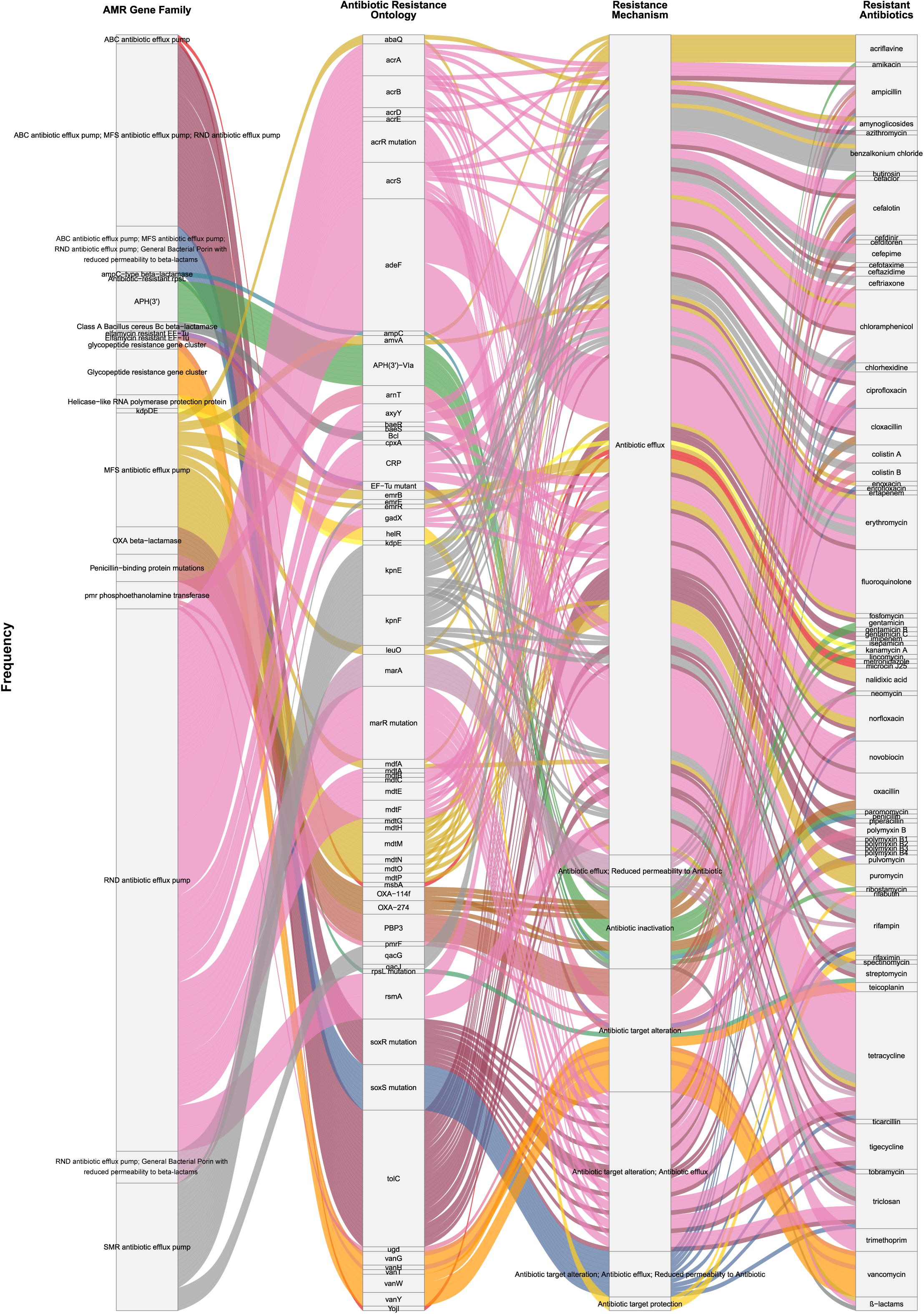
Resistome profile of the GPMAGs. Identified AMR gene frequencies in the GPMAGs were categorized by gene family, resistance mechanism, and target antibiotics.

**Fig. S3.**
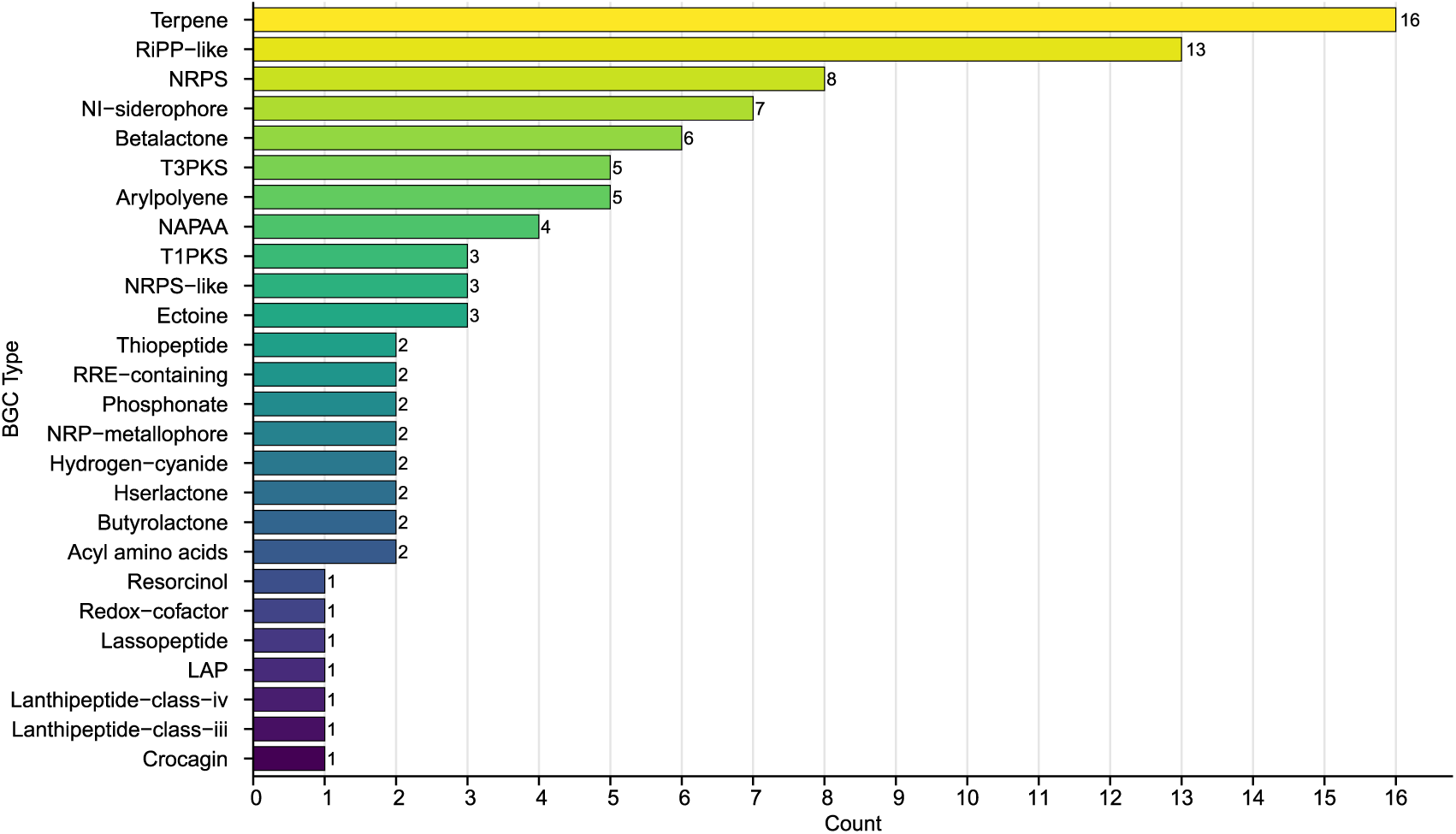
BGC profile of the GPMAGs. Identified BGCs from the GPMAGs were displayed by bar chart.

**Fig. S4.**
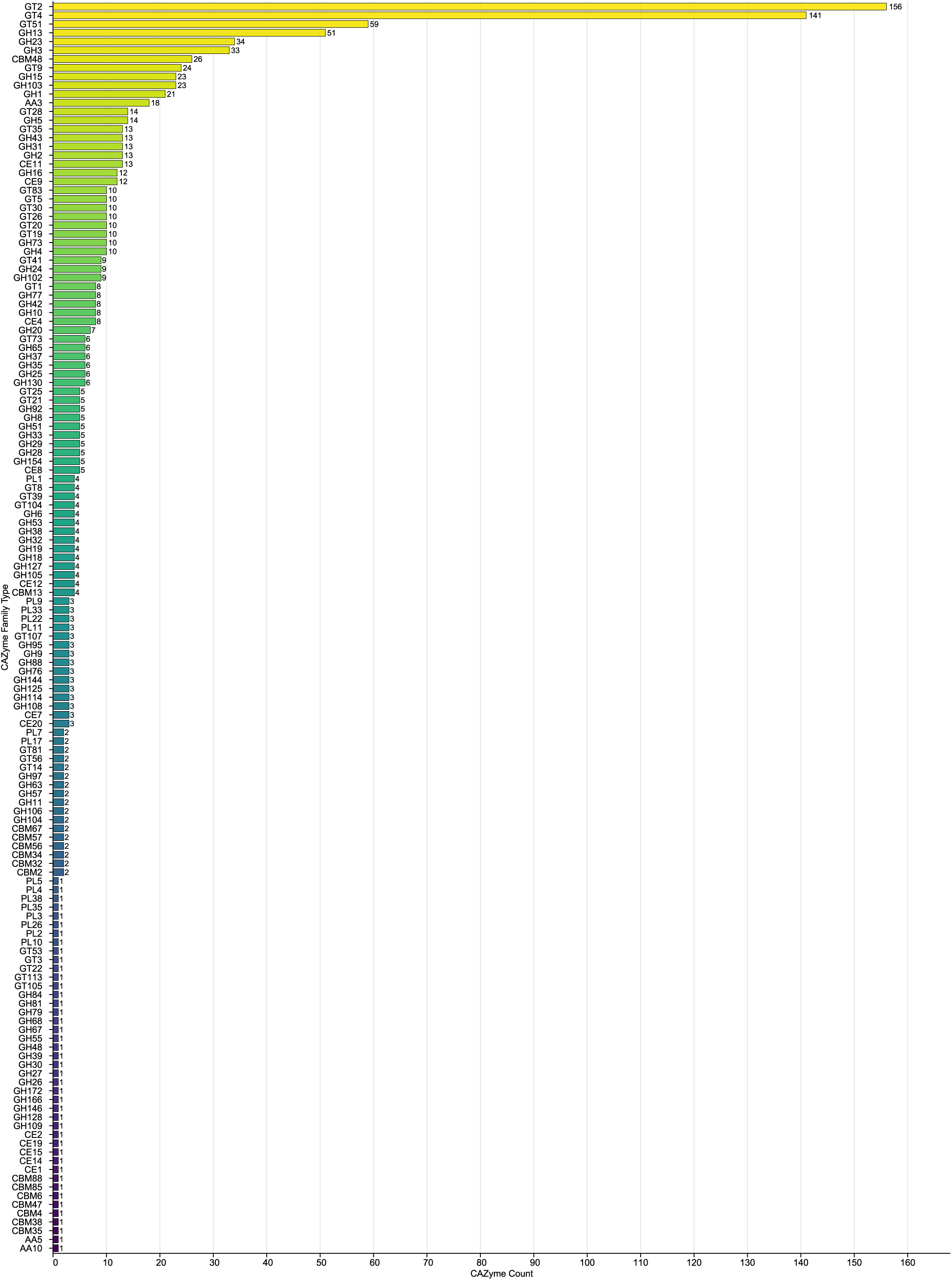
CAZyme profile of the GPMAGs. Bar chart exhibits identified CAZyme types of the GPMAGs.

**Fig. S5.**
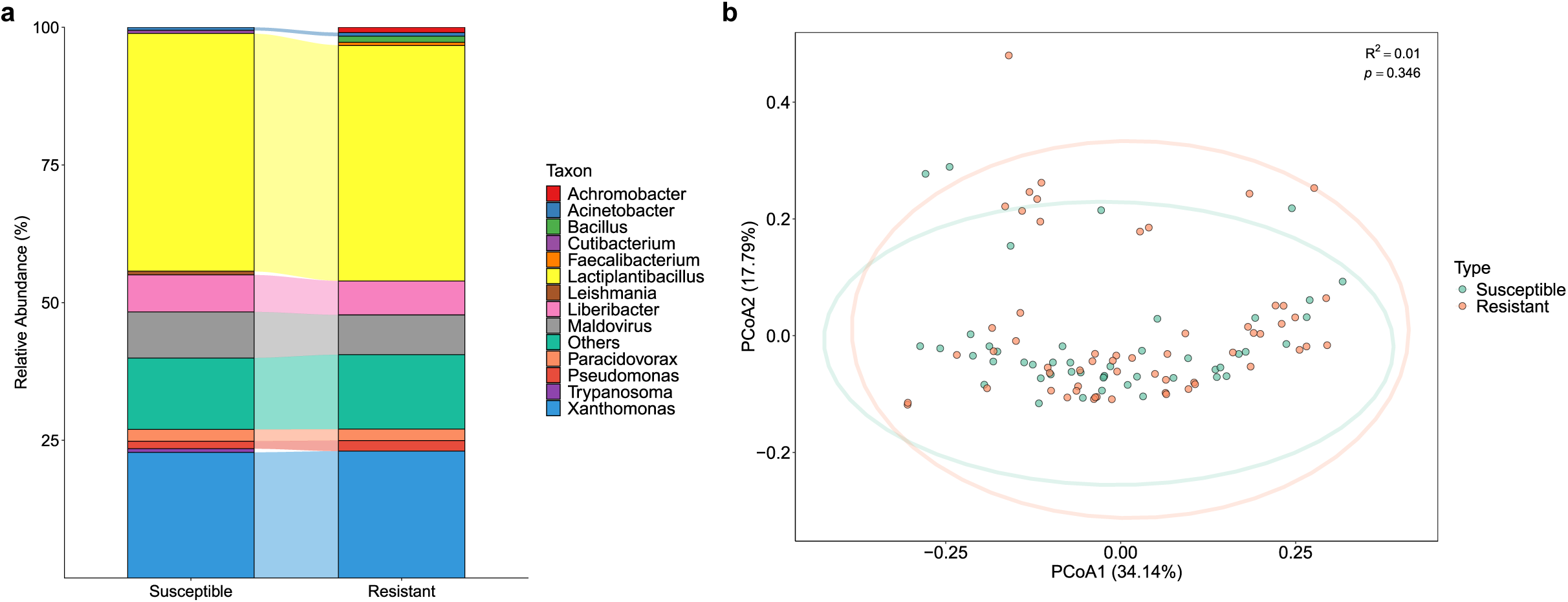
Microbiome profiles associated with DM resistance. **a**, Mean relative abundances of the top 10 most abundant genera based on the types of DM resistance. **b**, PCoA of the grape microbiomes at genus level. Each point represents a grapevine accession with colored according to its variety. Ellipses denote the 95% confidence intervals for each group. Percentages on the axes indicate the variance explained by PCoA1 and PCoA2. PERMANOVA analysis revealed significant differences among the groups, with R² values represent the proportion of variance explained by the grouping factor and p-values indicate the statistical significance of these differences.

**Fig. S6.**
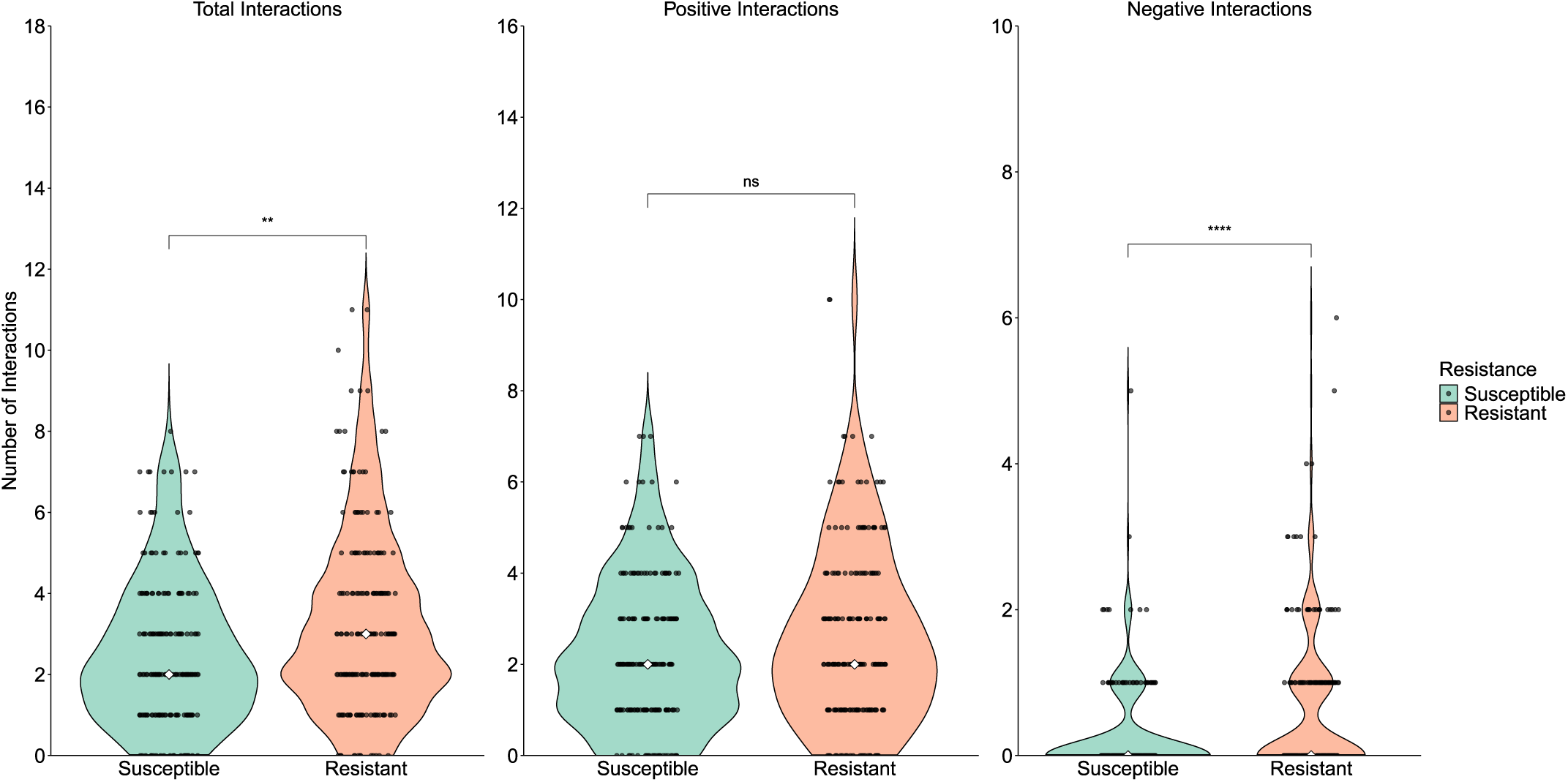
Microbial interaction profiles associated with DM-resistance. Violin plots depict the median microbial interactions for each node in the networks shown in Fig. 5a. White diamonds indicate medians. Significance levels are denoted as ** (*p* < 0.01), **** (*p* < 0.0001), and ns (not significant).

**Fig. S7.**
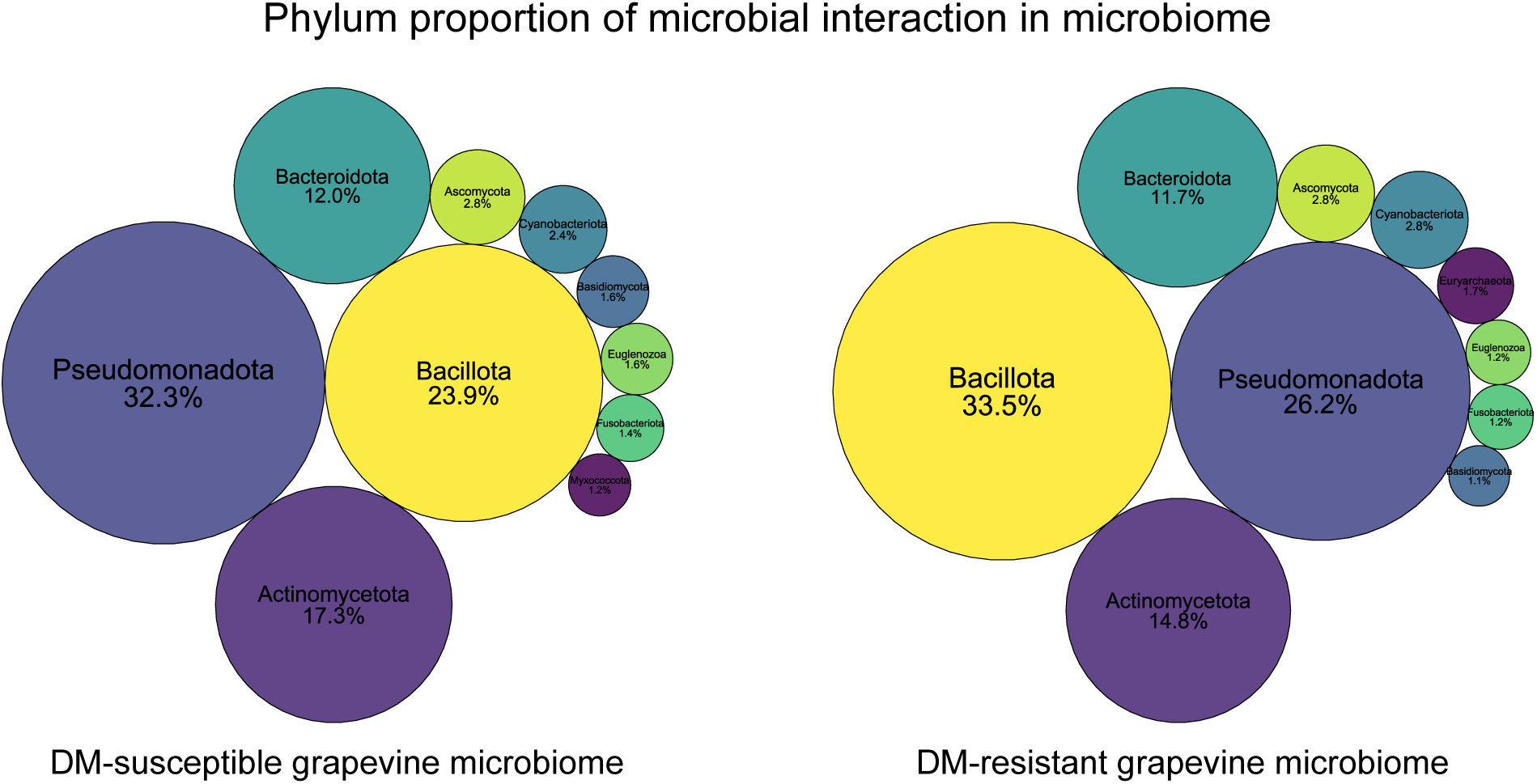
Phylum-level microbial interaction proportions between microbiomes. Bubble plots display interaction frequencies among the top 10 abundant phyla in the susceptible and resistant microbiomes, with bubble sizes corresponding to proportional values (numerically labeled).

**Fig. S8.**
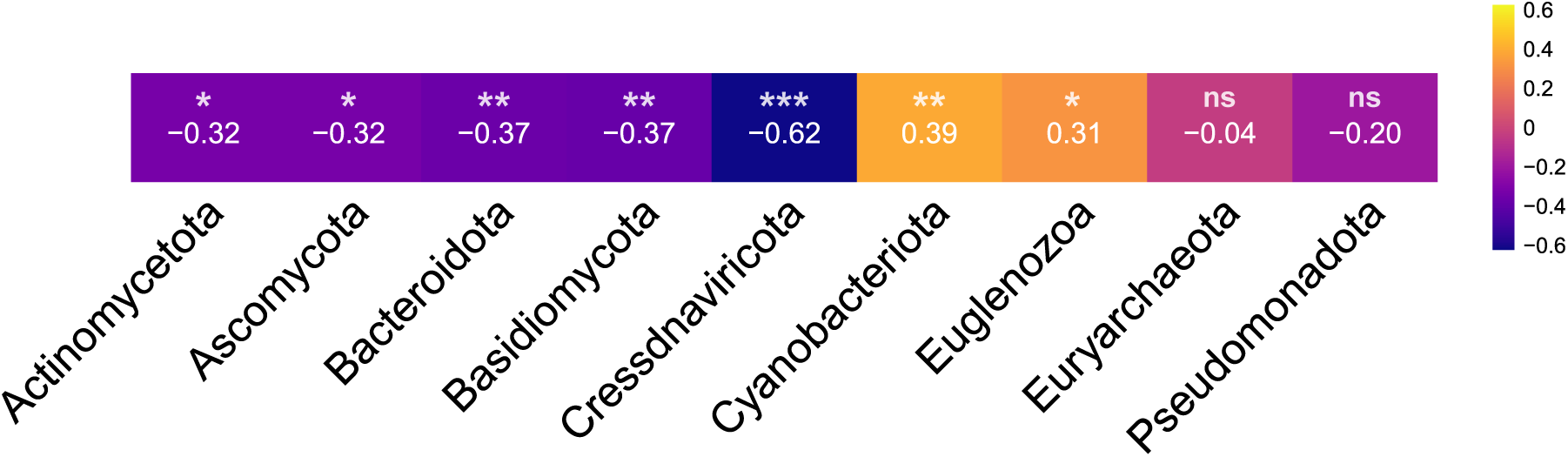
Abundance correlation between Bacillota and major phyla in the resistant microbiome. Heatmap depicts Spearman coefficients between Bacillota and major phyla based on their abundances in the resistant microbiome. Numerical values and significance levels * (*p* < 0.05), ** (*p* < 0.01), *** (*p* < 0.001), ns (not significant) are shown in cells.

